# Are experiment sample sizes adequate to detect biologically important interactions between multiple stressors?

**DOI:** 10.1101/2021.07.21.453207

**Authors:** Benjamin J. Burgess, Michelle C. Jackson, David J. Murrell

## Abstract

As most ecosystems are being challenged by multiple, co-occurring stressors, an important challenge is to understand and predict how stressors interact to affect biological responses. A popular approach is to design factorial experiments that measure biological responses to pairs of stressors and compare the observed response to a null model expectation. Unfortunately, we believe experiment sample sizes are inadequate to detect most non-null stressor interaction responses, greatly hindering progress. Determination of adequate sample size requires (i) knowledge of the detection ability of the inference method being used, and (ii) a consideration of the smallest biologically meaningful deviation from the null expectation. However, (i) has not been investigated and (ii) is yet to be discussed. Using both real and simulated data we show sample sizes typical of many experiments (<10) can only detect very large deviations from the additive null model, implying many important non-null stressor-pair interactions are being missed. We also highlight how only reporting statistically significant results at low samples sizes greatly overestimates the degree of non-additive stressor interactions. Computer code that simulates data under either additive or multiplicative null models is provided to estimate statistical power for user defined responses and sample sizes and we recommend this is used to aid experimental design and interpretation of results. We suspect that most experiments may require 20 or more replicates per treatment to have adequate power to detect non-additive. However, researchers still need to define the smallest interaction of interest, i.e. the lower limit for a biologically important interaction, which is likely to be system specific, meaning a general guide is unavailable. Sample sizes could potentially be increased by focussing on individual-level responses to multiple stressors, or by forming coordinated networks of researchers to repeat experiments in larger-scale studies. Our main analyses relate to the additive null model but we show similar problems occur for the multiplicative null model, and we encourage similar investigations into the statistical power of other null models and inference methods. Without knowledge of the detection abilities of the statistical tools at hand, or definition of the smallest meaningful interaction, we will undoubtedly continue to miss important ecosystem stressor interactions.

## Introduction

Most, if not all, ecosystems are being impacted by multiple co-occurring stressors (e.g., climate change, invasive species, pollution), which are predominately anthropogenic in origin (Halpern et al. 2015; Beauchesne et al. 2021), and are capable of affecting individuals through to entire ecosystems (Jackson et al. 2021; Simmons et al. 2021; Sokolova 2021). At the individual level, responses to multiple stressors might be assessed by their joint effect on the physiology of an organism, e.g., a decline in feeding, growth, or fecundity, or a biochemical change (Nõges et al. 2016), and may also be measured on survival rates (e.g. bee health responses to agrochemicals, Siviter et al. 2021). Population responses to multiple stressors may be assessed by monitoring densities, biomass, or other markers such as chlorophyl concentrations (e.g. freshwater population responses to combinations of invasive species, pesticides, temperature or UV changes, Burgess et al. 2021), whereas ecosystem responses might be measured through multiple stressor effects on functional and taxonomic diversity (e.g. coral reef species richness responses to warming and acidification, Timmers et al. 2021), or through other measures on ecosystem integrity (e.g. stability, Polazzo and Rico, 2021).

Going beyond effects of single stressors is therefore an important focus in ecology and a key question is whether and how these co-occurring stressors may interact. For example, two stressors operating together may act to amplify their individual effects and lead to a synergistic interaction. In this case their joint effects are greater than predicted from their individual effects. This might occur for example if one stressor (e.g. dehydration caused by a drought) reduces the fitness of an individual and makes it more susceptible to another stressor such as a disease (Lafferty and Holt, 2003). On the other hand, two stressors acting on the same biological process could have a negative (interfering) effect on one another and therefore lead to an antagonistic effect; their joint effects are less than predicted by their individual effects. In extreme cases this can lead to reversal interactions (Jackson et al., 2016) where the combined effect of a pair of stressors has a different sign to those of both stressors acting on their own. For example, Boone et al. (2005) showed how the combined effect of carbaryl and nitrate decreased green frog (*Rana clamitans*) tadpole growth, even though individually both increased tadpole growth.

Cataloguing, and predicting how often and under what conditions synergies and antagonisms might occur can have important implications for management strategy. In the case of a synergistic interaction between two stressors, removal or reduction of the impact of even one stressor could have a large effect. However, more caution is required when considering management of an antagonistic interaction since, if the antagonism is particularly strong, removal of one of the stressors could in principle lead to a worse outcome as the biological response to the pair of stressors might be less severe than the response to either stressor acting alone. However, current knowledge of how stressors interact to affect biodiversity at various scales is limited (Hodgson and Halpern 2019; Lemm et al. 2021). To date, progress has been driven by individual studies that have contributed to larger-scale meta-analyses, but relatively few generalisations are possible (Côté et al. 2016; Orr et al. 2020). This is perhaps not surprising given the broad range of ecosystems, taxonomic groups, and biological responses that have been considered (e.g., Ban et al., 2014; Burgess et al., 2021; Lange et al., 2018), but another contributory factor that has not been examined is the issue of adequate sample sizes in multiple stressor experiments.

We contest that many potentially important stressor-pair interactions are being missed due to low replication number. In order to design effective multiple stressor experiments that have adequate sample sizes, researchers must consider the trifecta of: i) resource costs (whether the design is feasible given time, spatial, financial constraints), ii) the smallest stressor-pair interaction that can be detected (statistical power), and iii) the minimum biological effect of interest (Figure 1). However, we believe only resource costs and therefore feasibility normally factor into experimental design since the detection limits of the statistical tools commonly used in stressor interactions have not been quantified, and there has been no discussion on what a biologically important stressor interaction is. We define the smallest interaction of interest as the smallest biologically relevant deviation from the null expectation and could represent the smallest deviation that would warrant a change in management strategy compared to the null. Here we will look at sample sizes typical of stressor interaction experiments, use empirical examples, and analyse of statistical models to highlight why it is likely important interactions are being missed, and show how the minimum biological effect of interest dictates the sample sizes required.

**Figure 1.**
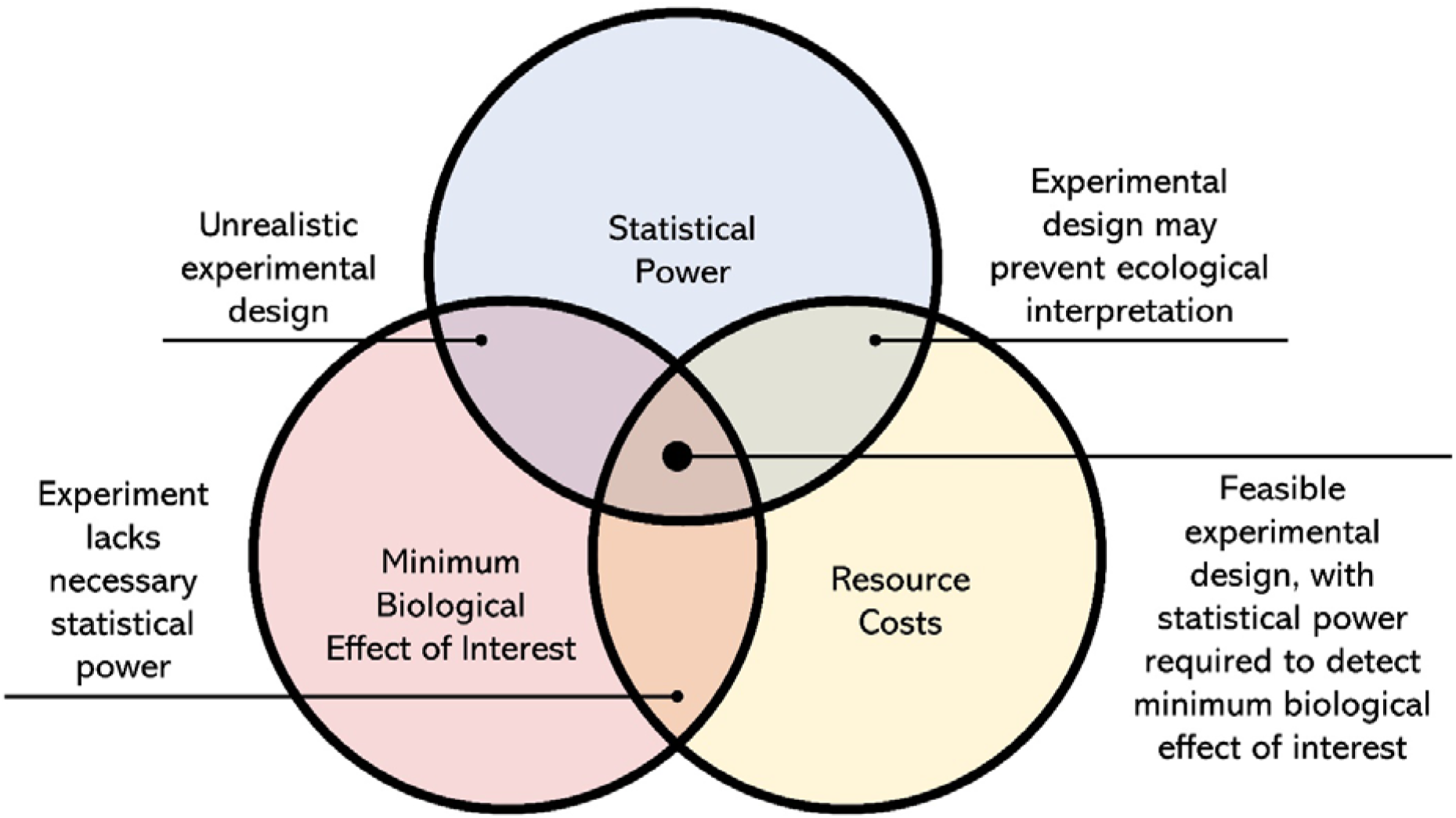
The three considerations important for determining experimental design to investigate how pairs of stressors interact, and the trade-offs that occur when any of them are more limiting than the others.

### Stressors: model expectations and interactions

The effects of multiple interacting stressors are commonly determined through the implementation of null models (e.g., Schäfer and Piggott, 2018) where the observed response is compared to an expectation that the stressors are non-interacting (De Laender, 2018). Other methods are available, such as the linear model approach (e.g., Spears et al. 2021), but null models continue to enjoy widespread use in ecology and evolution (e.g. van Veen and Murrell, 2005; Flügge et al. 2012; Murrell, 2018; Rajala et al. 2018), Moreover, linear models also make assumptions about the form of the interaction (e.g. additive) and in any case the issue of sample size is germane to all approaches. Of the range of available null models for multiple stressor interactions, the additive null model (Gurevitch et al., 2000) is the most widely applied (e.g., Crain et al., 2008; Burgess et al. 2021; Siviter et al. 2021) and has the expectation (null hypothesis) that the overall effect of the multiple interacting stressors is equal to the sum of the effects of the stressors acting individually. In effect the question is: “Do the individual effects of two stressors simply add up when they are both present?”.

The statistical test is therefore whether the additive null model can be rejected in favour of an alternative hypothesis that interactions are: i) greater than anticipated by the additive null model (*Synergistic interactions*); ii) less than the sum of the individual stressor effects (*Antagonistic interactions*); or iii) opposite to that suggested by the additive null model (*Reversal interactions*) (see e.g., Jackson et al., 2016; Orr et al. 2020). Although we will focus on the additive model and show it has low power to detect non-additive stressor-pair interactions, we also show similar results for the multiplicative null model (Lajeunesse, 2011), which is argued (Fournier et al., 2006), to be preferable for biological responses (e.g., survival) that are bounded (see Supporting Information).

The null model approach requires a factorial experiment design with four treatments that each measure the same biological response metric of interest (e.g., individual survival; population density or biomass; species richness) under different stressor conditions. Each measure 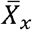, is the mean value of this response metric taken over *N*_*x*_ replicates, where *x* ∈ {*C, A, B, I*}. The first treatment, *C*, is the control which is the system (i.e., individual, population, community) of interest in the absence of either stressor under scrutiny. There are two treatments (A, B) that account for the response of the system to each of the individual stressors of interest acting in isolation. The final treatment, *I*, is the estimate of the response to both stressors acting simultaneously i.e. the interaction. Associated with each treatment is an estimate of the standard deviation of the response to the treatment, and these are denoted by *SD*_*x*_, where again *x* ∈ {*C, A, B, I*}. All three elements, 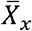, *SD*_*x*_, and *N*_*x*_ are required for the additive and multiplicative null models and from this input each null model computes an effect size, with associated confidence intervals from which the interaction type is inferred.

Effect sizes are used as they can provide a standardised measure of the difference between two groups (treatments) and therefore enable straightforward comparison of experiments where the biological response may be on different scales (e.g. density, survival). In the case of stressor-pair interactions the effect size is defined as the difference between the response predicted by the null model from the individual responses (A and B) and the observed response to both stressors acting simultaneously (I). We use the definition of effect sizes for factorial experiments under the additive model defined by Gurevitch et al., (2000). The observed interaction effect is defined as 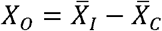, and the expected response that assumes the joint effect is equal to the sum of the individual effects of stressors *A* and *B* is defined as 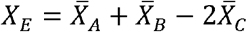. To compute effect sizes (*ES*_*Add*_), we use Hedges’ d which is unbiased by small sample sizes (Hedges and Olkin, 1985). The calculation of the additive effect size, (*ES*_*Add*_), is given as

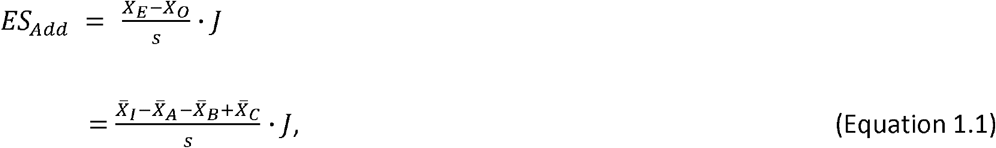

where *s* is the pooled standard deviation that takes into account the standard deviations (*SD*_*X*_) associated with each treatment mean, and *J* is the small sample bias correction factor (Borenstein et al., 2009). Both s and *J* are defined in the Supporting Information.

Once computed, we need to know if *ES*_*Add*_ is statistically different from 0 in which case the nullvhypothesis is rejected in favour of an alternative that is dependent on whether *ES*_*Add*_ is positive or negative (explored in more detail in the Supporting Information). Put simply, the test answers whether there is sufficient evidence to define the stressor interaction as being non-additive. The test requires the construction of confidence intervals (at some specified level of statistical significance *α*), and these in turn require an estimate of the standard error for our effect size. The estimate of the variance defined by

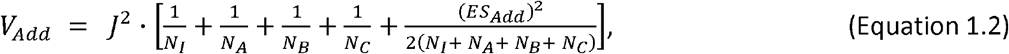

and from this the standard error is computed as

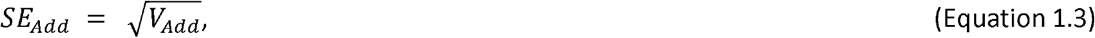

with the important observation that the standard error for *SE*_*Add*_ is not divided by the square root of the sample size as is the case for normal estimates of the sampling distribution of a mean. Standard errors should decrease as more samples are taken but increasing sample sizes will already reduce the variance (Equation 1.2), and hence *SE*_*Add*_ Finally, the confidence intervals are computed as

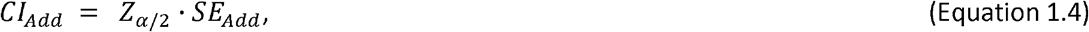

with *Z*_*α*/2_ being the critical Z-score taken at the statistical level of significance *α* Typically, *α* = 0.05, and we divide by two as a two-tailed test is required because the stressors interaction can be less than, or greater than expected under the null model, which means *Z*_*α*/2_ = 1.96. The test has *df* = *N*_*I*_ + *N*_*A*_ + *N*_*B*_ + *N*_*C*_ − 4 degrees of freedom. An important point to note is how the sample sizes *N*_*x*_ appear at multiple stages in the process, with increasing sample sizes leading to smaller confidence intervals for the effect size, and a higher chance that the null hypothesis is rejected (because 0 is not contained within the range covered by the confidence intervals). As the equations contain many terms, it is relatively easy for a small error to creep into the computation of the effect sizes and confidence intervals, although this may be avoided through the use of openly available statistical software such as the R library *multiplestressR* (Burgess and Murrell, 2021).

In case the reader is in any doubt about the potential importance of interactions relative to the single stressor effects we use data on bee responses to a range of agrochemicals, nutrient stressors and parasites published in Siviter *et al*. (2021) to highlight how single stressor and multiple stressor effect sizes have similar overall distributions (Figure 2). What is also clear is that, at least in this data, interaction effect sizes may be quite large even though single effects are negligible and vice versa. Therefore, absence of large effect sizes in biological responses to individual stressors does not preclude the possibility for large effect sizes for the interaction, i.e. the interaction may be very different to the null expectation (and therefore non-additive) even though responses to individual effects are negligible.

**Figure 2.**
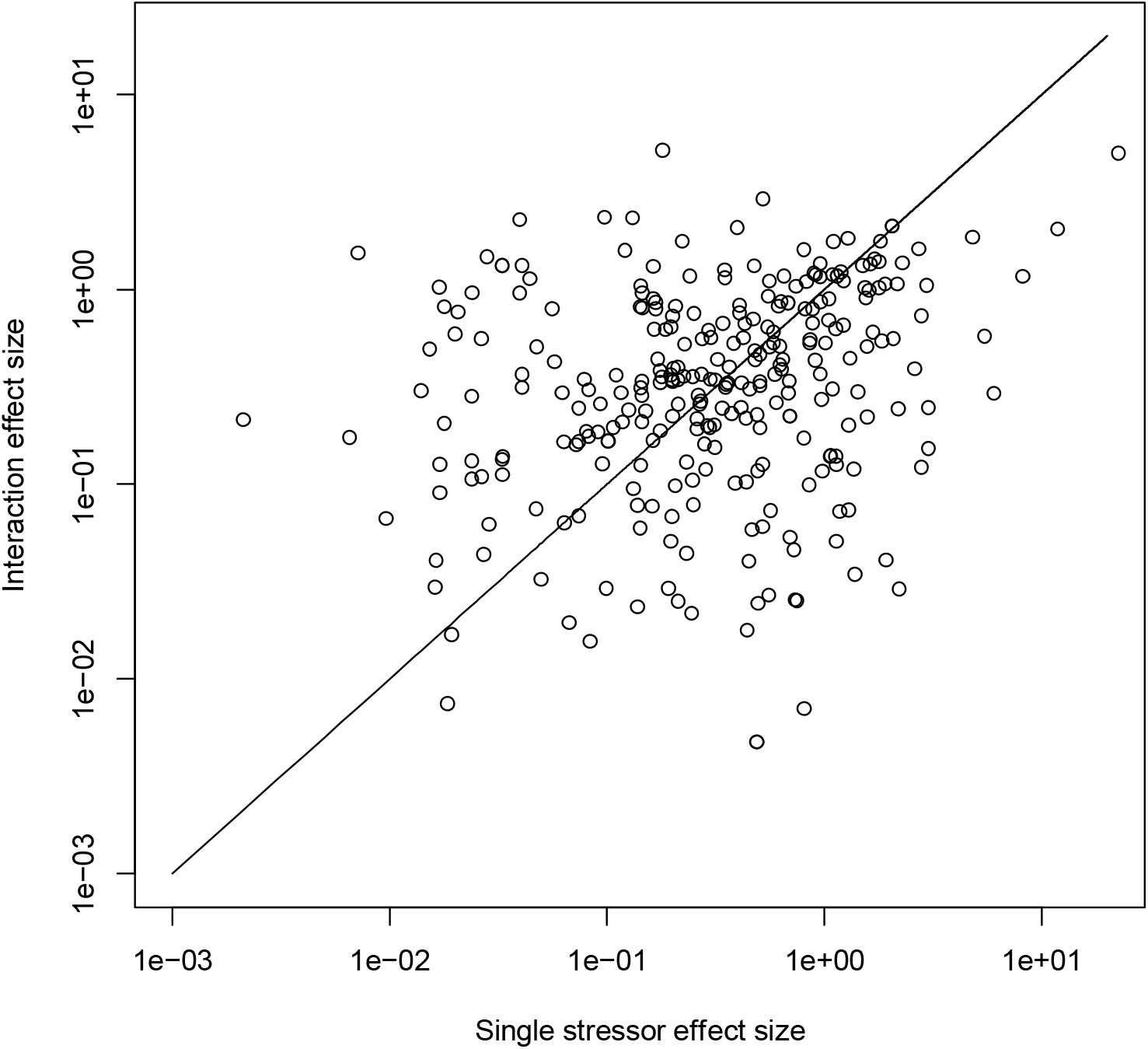
Scatter plot of Hedge’s d effect sizes for bee health response to single stressors (x-axis) and the interaction of two stressors (y-axis). Data is taken from the meta-analysis of Siviter et al. (2021), and we plot the absolute value for the effect sizes on a logarithmic scale. Interaction effect sizes (*ES*_*ADD*_) are computed assuming the additive null model, using equation (1.1). Single stressor effect size is computed using the *escalc* function in the R library *metafor* (Viechtbauer, 2010). The straight line is the line *y* = *x*, therefore denoting the special case where the absolute value of the single and interaction effect sizes are equal. Points below this line denote single stressor effect sizes larger in absolute value than stressor pair interaction effect sizes and those above the line denote the opposite relationship.

### Typical samples sizes in multiple stressor experiments

Perhaps the most basic question an empirical scientist can ask is “Does my study have sufficient data to answer my question?” (Johnson et al., 2015). In multiple stressor research this amounts to asking whether the sample size is sufficient to detect a departure from the null model of a *given magnitude* should this be the true interaction. We emphasise the qualification of a *given magnitude* as this is where the researcher has to determine *a priori* the smallest deviation from the null expectation that is biologically important. However, this concept has not been discussed, but is critical to knowing how likely we are to be missing important non-null stressor interactions and is a point we focus on in more detail below.

In the absence of any guidance based upon understanding of the null models, researchers have to make sample size decisions that are likely more determined by resource constraints (financial, time, or space costs; Boyd et al., 2018; Rineau et al., 2019), or heuristic arguments (such as a rule of thumb value that is not based on power analyses). Perhaps as a consequence of the lack of statistical guidance, the number of replicates in experiments to investigate stressor interactions rarely reaches double figures. For example, two recent meta-analyses (Gomez Isaza et al., 2020; Seifert et al., 2020) included no experiments with more than six replicates per treatment, while a third (Burgess et al., 2021) found <1% of the experiments used more than eight replicates per treatment (Figure 3). Exceptions to this trend tend to focus on individual-level responses with recent examples taken from honeybee health responses to multiple pesticides (Bird et al. 2021) where the control treatment mean sample size was 179.33, and bee responses to pairs of agrochemicals where the control treatment mean sample size for studies where this data is publicly available was 115.62 (Siviter et al. 2021).

**Figure 3.**
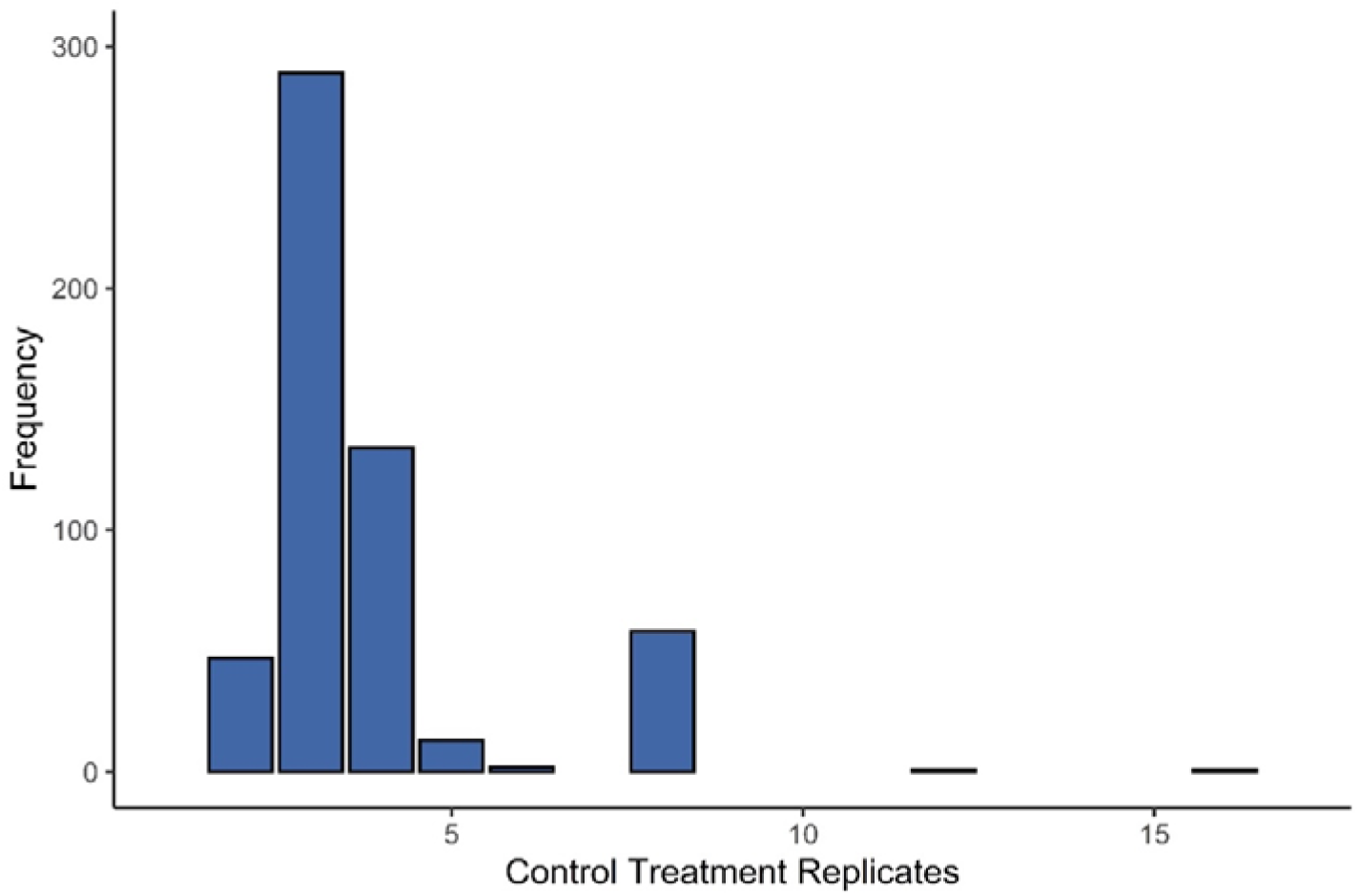
The frequency distribution of control treatment sample sizes from a dataset of 545 stressor interactions in freshwater ecosystems (Burgess et al., 2021).

The importance of sample size for detecting interactions between pairs of co-occurring stressors has only recently been acknowledged. Using simulated data created from a food web model Burgess et al. (2021) showed how even low levels of observation error, where 99% of all measured responses were within 10% of the true response value, can lead to the inability to detect the true, non-additive interaction in the majority of cases at typical sample sizes of *N*_*x*_ = 4. In other words, even small levels of noise can overwhelm the biological signal when sample sizes are low. Burgess et al. (2021) concluded that the large proportion of perceived additive interactions in their freshwater-focussed dataset could easily be explained by the low sample sizes (Figure 3), and that many possibly biologically important non-additive stressor interactions were being missed. However, whilst this warning is useful, it does not answer the question of how many replicates are required.

### Critical effect sizes: the smallest detectable interactions

The ability to detect a non-null interaction is dependent on the strength of the interaction, the variation of the biological responses, and the sample sizes (i.e., 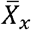, *SD*_*x*_, and *N*_*x*_), as well as the level of statistical significance *α*. Both 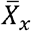 and *SD*_*x*_, are unknowns and are to be estimated in the experiments, whereas *N*_*x*_ (barring resource costs), and *α* are both choices of the researchers. The importance of sample size in detecting non-null interactions can be illustrated with an empirical example (Figure 4). Here, we use the additive null model to determine the effect of stressor pairs on bee health data (Siviter et al., 2021) which comprises a wide range of sample sizes. As expected, increasing sample size results in an increased ability to detect non-null interactions, and we can see how greater sample sizes allow weaker non-null interactions to be identified and classified (Figure 4).

**Figure 4.**
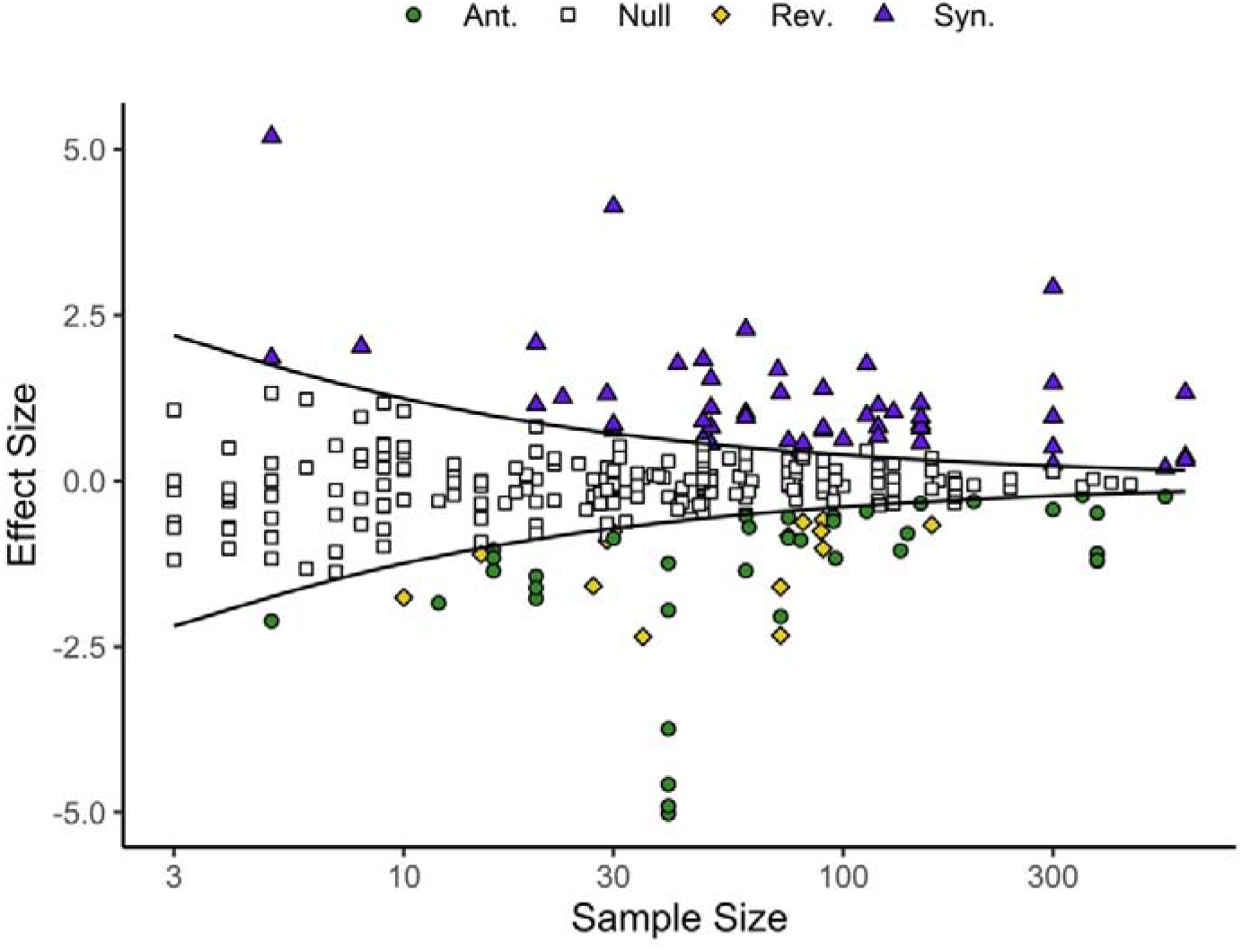
The effect of sample size on the ability to detect interactions with different effect sizes for the bee health responses to multiple stressors in Siviter et al. (2021). Open squares denote data points that are statistically indistinguishable from the null model of an additive interaction (i.e., the null model that co-occurring stressors are simply the sum of their individual effects). Data points that lead to the rejection of the null model can be assigned as synergistic (purple triangles), antagonistic (green circles), or reversals (yellow diamonds). The black lines denote the critical effect size that separates the region of detectable departure from the null model at the 5% level of significance. Median sample size per treatment is plotted on the x axis. A small number of null interactions appear outside of the null region where the experiment had uneven sample sizes between treatments, but for clarity of presentation the critical effect size is computed under the assumption of equal sample sizes within each study. Results were generated using the *multiplestressR* R package (Burgess and Murrell, 2021, 2022), with code to reproduce this figure provided in the Supporting Material.

For each sample size, there is a minimum effect size that an experiment will be able to distinguish as being statistically different to the null model (illustrated by the black lines in Figure 3). Effect sizes below this threshold denote interactions that cannot be distinguished from the null model expectation of additivity at the chosen level of statistical significance. This threshold, referred to as the *Critical Effect Size* (see Mudge et al., 2012; Lakens, 2022) can be exactly calculated for the additive null model (the equation for which is detailed in the Supporting Information but can be computed using the R library *multiplestressR*; Burgess and Murrell, 2021). Analysis of the bee health data (Siviter et al., 2021) shows how the critical effect size (*ES*_*Add*_) predicts non-additive interactions and verifies the expectation that only very large effect sizes can reject the null expectation of additivity when sample sizes are below 20 per treatment (Figure 4). At the very low samples sizes that typify multiple stressor research, especially for population- and community-level responses, effect sizes have to be very large (e.g., for *N*_*x*_ = 4, *ES*_*Add*_ ∼2) in order for non-additive interactions to be detected.

### Statistical power

The critical effect size is the smallest detectable effect size for a given sample size, but due to sampling variation we can expect the estimated effect size to differ between repeat experiments. Statistical power represents the proportion of these repeat experiments that would correctly result in the rejection of the null model expectation, assuming a non-additive interaction exists, and we explore this using a data simulation approach. Although any single effect size can be generated by an infinite number of combinations of treatment means and treatment standard deviations, we use a simple example to illustrate low sample sizes yield low power to detect non-additive interactions.

We set the expected control treatment mean biological response (e.g., survival probability) to 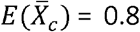. The expected responses to two separate stressors (e.g. pesticides, A and B) are assumed to be the same, and we set 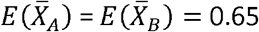, whereas the expected mean of the response to both stressors acting simultaneously is allowed to vary 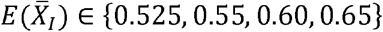. In all treatments the expected standard deviation *E*(*SD*_*x*_) = 0.05. These values for 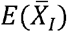 and *E*(*SD*_*x*_) gives rise to expected effect sizes *E*(*ES*_*ADD*_) = {3, 2, 1, 0.5} respectively. In all cases the interactions are less than the additive prediction and should result in an antagonistic interaction being inferred. For simplicity we assume all treatments have the same replication number, so *N*_*C*_ = *N*_*A*_ = *N*_*B*_ = *N*_*I*_ = *n*. We simulate 1000 ‘experiments’ for each combination of *n* and 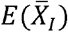, and assume treatment values are sampled from a Gaussian distribution with standard deviation *α*_*x*_ = *E*(*SD*_*x*_), and means given by the expected treatment means 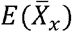, We then use *multiplestressR* (Burgess and Murrell 2021, 2022) to test whether we can correctly reject the null model of an additive interaction in favour of an antagonistic interaction for each ‘experiment’, and from this we compute the statistical power.

Simulating effect sizes under these parameters shows clearly that low sample sizes lead to low statistical power size (Figure 5a). For example, when *n* = 3, only about 50% of experiments would result in the correct rejection of the null model when the expected effect size is 3. The problems are predictably worse for smaller effect sizes, and even *n* = 20 results in power of only approximately 0.5 when the expected effect size is 1. To get power of at least 0.8 requires samples sizes of approximately 5, 9, 34 and >100 for *E*(*ES*_*ADD*_) = {3, 2, 1, 0.5} respectively. As shown in Figure 2, most empirical interaction effect sizes are below 1, and this means *n* > 18 is required to correctly reject the additive null model at least half the time. Adjusting the parameters to get the same effect sizes but with *α*_*x*_ = 0.025, for *x* ∈ {*C, A, B, I*} shows treatment variance makes a negligible difference (see Figure S2, Supporting information) and verifies earlier work that shows Gaussian distributed observation errors have to be unrealistically small (*σ*_*x*_ < 0.0001) in order to lead to a high detection rate (Burgess et al., 2021). However, as shown by Burgess *et al*. (2021) for *n* = 4, reducing treatment variation (i.e. lowering *E*(*SD*_*x*_) whilst keeping expected treatment means constant) will result in larger effect sizes and will therefore increase power to detect.

**Figure 5.**
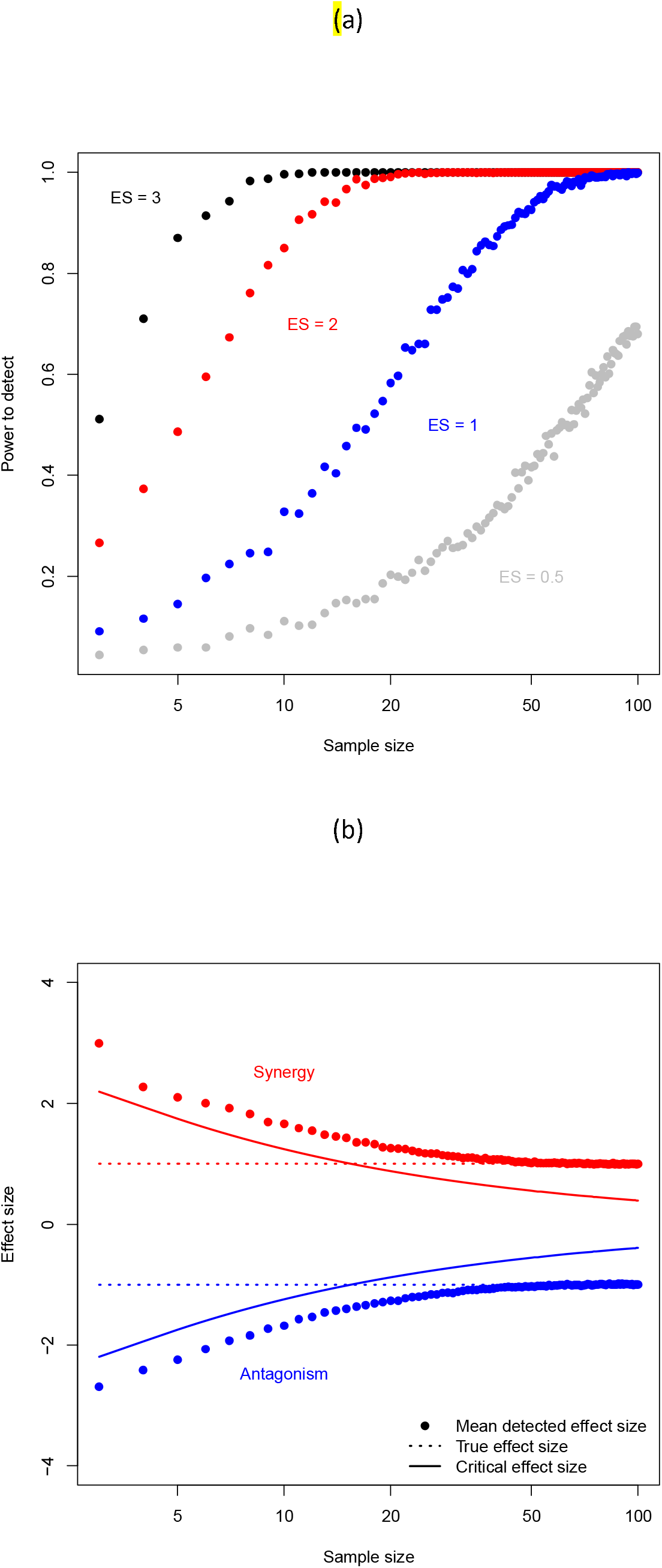
The effect of sample size on (a) the power to detect non-additive interactions of different strengths as determined by the effect sizes (ES); and (b) the bias towards overestimating the strength of the departure from additivity when considering only those interactions that result in a statistically significant result. Data is simulated with two stressors causing the same response when operating in isolation and all treatment standard deviations are set to have the same value. In (a) the expected interaction treatment mean is varied to generate the different expected effect sizes. In (b) the mean detected effect size averages over only those simulations where the null model is rejected. In both panels the data points are computed from 1000 simulations (‘experiments’) for the same set of parameters at each sample size. See main text for more details of the simulations.

A consequence of low statistical power is that considering only the statistically significant interactions may greatly overestimate the effect size and hence overestimate the deviation of the interaction from additivity. Figure 5b shows examples for a synergistic interaction 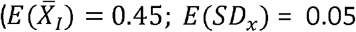, other parameters as before) and an antagonistic interaction 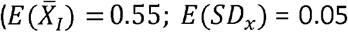, other parameters as before) for a range of sample sizes. The expected (or true) effect sizes are *E*(*ES*_*ADD*_) = 1, and *E*(*ES*_*ADD*_) = −1, respectively, The critical effect size determines the smallest effect size that can result in a non-additive interaction being detected, so detected effect sizes are always larger than this value. In our examples the mean detected interaction effect size only approaches the true interaction effect size at around *n* = 40, and at small sample sizes the mean detected effect size is approximately three times the magnitude of the true effect size (Figure 5b). This shows how publishing only statistically significant results from experiments with low sample sizes leads to overestimation of non-additivity, a problem that has also been highlighted for biological responses to single stressors (Yang et al. 2022).

### Smallest interaction of interest: *What is a biologically meaningful interaction?*

Up to now our discussion has largely related to *statistical* but *not biological* significance i.e. we have asked: (1) what is the smallest effect size we can detect, and (2) what is our statistical power for given sample size? As we have shown, small sample sizes can lead to the detection of only large effect sizes and therefore highly non-additive interactions (Figure 4), but at the other end of the scale infinitely large sample sizes can detect infinitely small departures from additivity (i.e., the lines in Figure 4 asymptote slowly to 0). So, whilst small sample sizes likely miss key stressor interactions, large sample sizes can waste resources (Figure 1) and uncover biologically insignificant stressor-pair interactions. To avoid either of these outcomes, the researcher needs to determine the smallest interaction that would lead to a biologically meaningful deviation from the null model before the experiment is run (to avoid any bias from knowing the result). We define this interaction as the minimum biological effect size, and we argue this depends upon both the study system and response of interest. For example, a researcher may want to determine whether two stressors combine to affect a response (e.g., juvenile survival rates) in a non-additive manner for an endemic or threatened species. In this scenario it is important to be able to detect a small deviation from additivity (i.e., a small effect size) as failing to detect even a weak interaction may lead to the wrong mitigation strategy being selected and potentially exacerbate the effects of these stressors to the detriment of the study system (Brown et al., 2013; Côté et al., 2016). Commonplace sample sizes (e.g. 4 replicates per treatment) are not adequate for this question (Figures 4, 5), and the researcher will likely need to implement sample sizes that are multiple (two or more) times larger than those commonly used. There may be other situations where a smaller effect is not so important, implying smaller samples are adequate, such as monitoring abundance declines in a system with high functional redundancy, but even here care needs to be taken since concerns have been raised regarding publication bias leading to the overestimation of stressor effects from experiments with small sample sizes (Figure 5b, Yang et al., 2021).

How should the minimum effect size of interest be determined? Although it might seem tempting to use the heuristic guidelines proposed by Cohen (1988) for small, medium, large effect sizes, we do not believe they are appropriate for multiple stressor research due to the heterogeneity in systems, responses, and stressors. For example, would we decide upon the same minimum effect size for survival responses at different stages in a species’ life cycle? In any case these guidelines only relate to Cohen’s *d* or Hedge’s *d* and do not apply to null models such as the multiplicative null model that operate on a different scale. Other ways that the minimum effect size of biological interest could be determined include guidance from ecological theory, and results of previous meta-analyses (Lakens, 2022). However, in order for a theoretical model to be a useful guide, it needs to be an adequate approximation to the stressors, biological system, and response under scrutiny. This is a tall ask, since it is likely that empirical evidence is required to calibrate the model in the first place, in which case there is already some evidence that could be used (carefully) to consider the number of replicates required. The results of previous meta-analyses could act as a guide, although again care needs to be taken since it is possible that publication biases towards biologically novel but not necessarily statistically robust effect sizes (Filazzola and Cahill Jr. 2021) could affect summary effect sizes. Moreover, meta-analyses in ecology and evolution often report high levels of heterogeneity (Senior et al., 2016) compared to human clinical trials since ecological and evolutionary studies often focus on multiple taxa, in real world environments, that are subject to many different forms of environmental and biological variation (Burgess et al. 2021; Côté et al., 2016). It is therefore hard to know if the summary effect sizes reported in these meta-analyses are relevant for other, more focussed, studies that might being asking subtly different questions involving, for example, different stressors or responses.

## Consequences and recommendations

Overall, we do not believe there is a simple answer to the smallest effect size of biological interest. Instead, we propose researchers use their expert knowledge to use values for the treatment means and standard deviations and estimate power using the simple R function (*interaction_power*) we used to generate Figure 5. For example, it might be decided that a 10% deviation from additivity would constitute a biologically important stressor interaction, and along with estimates of treatment means and standard deviations the code could be used to explore likely levels of statistical power for a range of sample sizes. This will give at least a ball-park figure before the experiment is completed and may give the opportunity to increase sample sizes as appropriate. We also add that the code can be employed to estimate power for either additive or multiplicative null models (see Supporting Information). More generally, the sweet-spot of sample size is dependent on the trifecta of resource costs, statistical power, and minimum effect of biological interest, and failure to take any of these into consideration may limit the effectiveness of any experiment (Figure 1). However, it seems likely that in many cases *N*_*x*_ = 4 does in fact lead to biologically important on-null stressor-pair interactions being left undetected (Figures 4 and 5), and given the relationship between critical effect size and sample size, 20 replicates (or more) might be desirable.

The recent meta-analyses of how pairs of pesticides interact to affect bee health (Siviter et al. 2021, Bird et al. 2021) are examples of experiments with very large sample sizes, and the fact that they both focus on studies at the individual-level highlight how this might be a resource efficient way to increase replicate numbers. This echoes earlier calls to focus on individual-level responses to stressors as it is the fate and/or behaviour of the individual that is directly affected (e.g., Maltby 1999). However, responses at other (higher) levels of biological complexity such as population, community and ecosystem are also likely to be of interest because it is the response of these levels that may matter the most from a stressor management standpoint (Simmons et al. 2021). Moreover, because each species is embedded within a food web, interactions between species can lead to compensatory (antagonistic) or synergistic effects that are not observed for individual species in isolation (Christensen et al., 2006; Burgess et al. 2021; Simmons et al. 2021). Unfortunately, it is much harder to increase the sample sizes of many mesocosm experiments for these higher levels of organisation simply due to the financial cost, space, and time required to manage large sample sizes for all four treatments (Boyd et al., 2018). One alternative to boost within-study replication is to use coordinated networks of researchers who ask the same experimental question(s) across multiple sites, using the same protocol (Filazzola and Cahill Jr. 2021; Yang et al., 2022). An example of this is the Nutrient Network (NutNet) organisation (https://nutnet.org/) that amongst its key questions asks: To what extent are plant production and diversity co-limited by multiple nutrients in herbaceous-dominated communities? Another instance of this linked approach is the Managing Aquatic ecosystems and water resources under multiple stress (MARS) project (Hering et al., 2015) that has investigated the responses of a large number of European water bodies to multiple stressors (e.g., Birk et al. 2020). As always, there is no silver bullet, and coordinated networks may suffer from increases in data heterogeneity due to the multiple site nature of the network and the natural environmental and biological variation this includes, but also because small, but important differences in protocol may occur simply due to the number of research teams implementing the framework (Filazzola and Cahill Jr. 2021).

Our discussions of null models and sample sizes have been restricted to investigations of pairs of stressors, yet we know that many ecosystems are being challenged with more than two stressors (Halpern et al., 2015). For example, Nõges et al. (2016) identified European waters with up to seven co-acting stressors, although two co-acting stressors were the most common, being identified in 42% of cases. Similarly, there have been calls for investigating the responses to stressors at multiple levels of intensity (Polazzo et al. 2021; Schäfer and Piggott, 2018), since responses at low and high stressor intensities may differ greatly (Beaumelle et al., 2020; Dixon et al., 2020) and result in different interactions being detected (Ma et al., 2020). In both cases, sample sizes will need to be even larger than for two stressors each at a single intensity, and as we have already found, many experiments are probably greatly underpowered even in this simpler scenario. In order to maximise the outcome for the input of resources we suggest that individual studies should first try to boost sample sizes for simpler experiments before adding in further complexity, and encourage investigations of greater than two stressors and/or multiple intensities to use coordinated networks where the sample sizes can be distributed across multiple research teams, or focus on individual-level responses where sample sizes may more easily run into the hundreds (e.g. Bird et al. 2021; Siviter et al. 2021).

Ultimately, resource constraints may mean it is not possible to design an experiment with adequate sample sizes to capture biologically interesting/important stressor-pair interactions, especially for studies on responses at higher levels of biological organisation. Interpretation of experiments based on low sample sizes should be cautious and it should be remembered that failure to reject the null model is not evidence that mean that the null model is true. Hence, failure to detect a non-additive interaction between two stressors should not be associated with conclusions that the interaction is additive, only that there is insufficient evidence to show otherwise. Alternative statistical tests such equivalence tests (Lakens, 2017) are required to determine if any deviation from the null expectation is trivially small, and that the interaction can therefore be deemed additive. However, experiments with small samples are useful as they can provide data for meta-analyses that collate individual experiments together to greatly increase the power to correctly reject the null model (e.g., Crain et al., 2008; Jackson et al., 2016; Przeslawski et al., 2015). The key point is that to aid general understanding, and avoid publication bias (e.g. Figure 5b), it is crucial that all experiments are published with the data made openly available (i.e., the three components of sample size, mean and standard deviation/error or variance for each treatment) and not just those experiments that detect ‘interesting’ non-null stressor-pair interactions (Filazzola and Cahill Jr., 2021). Indeed, it is likely that publication bias is leading to the effects of anthropogenic stressors being overestimated (Yang et al., 2022), while multiple stressor ecology suffers from the erroneous over-reporting of synergistic interactions (Côté et al., 2016). Unfortunately, there are still many papers that do not report or make their data (i.e., treatment means etc.) readily available. For example, Burgess et al. (2021) identified 122 papers that appeared suitable for their meta-analysis of freshwater stressor interactions, but 66 had to be discarded due to missing data or having figures that were too unclear for data extraction. Not reporting these data represents a waste of resources, as it prevents future analyses (which are often unanticipated during the original study) from being conducted (Hanson and Walker, 2020).

In summary we make two main recommendations. Firstly, we urge researchers to make all data (sample sizes, mean and standard deviation of each treatment) easily available, regardless of statistical significance. Secondly, we ask researchers to state observed effect size(s), the critical effect size(s) if using the additive null model, and give an estimate of statistical power (e.g., by using data simulated using our code) of the experiment(s). Giving all this extra information will help to give an idea of the adequacy of the sample size implemented, and will also aid interpretation of the results.

## Conclusions

Our aim here was to open the discussion regarding sample sizes in multiple stressor research and show that before we ask the question “how much data do I need?”, we first need to answer the question “what is a biologically important interaction?”. Increasing sample sizes will always lead to an improvement in our statistical ability to detect unexpected stressor-pair interactions, but at extreme sample sizes we will likely be detecting only very small departures from the null model and these may not necessarily be relevant for management decisions. Setting the lower bound for an interesting stressor-pair interaction is critical to knowing what sample sizes are required. This lower bound is very much dependent on the system, stressors and response variable being measured, so we believe it can only be tackled using expert knowledge. Currently, it is our view that many experiments are likely underpowered and missing biologically important interactions, but studies that mostly focus on individual-level responses to stressors may be more adequately sampled. Strategies such as research networks may help increase sample sizes for higher levels of biological organisation such as communities, but there is still value in conducting smaller-scale studies, provided they are all published to avoid publication bias, and the data is made freely available, since they can contribute to meta-analyses and aid the design of subsequent experiments. We also urge the reporting of estimated power which will aid interpretation of results. Finally, although we have focussed on the commonly used additive and multiplicative null models, there are a number of other null models that have been proposed (e.g., Schäfer and Piggott, 2018; Dey and Koops, 2021), and to date there is no guidance on sample sizes required to detect non-null interactions of any given magnitude. This needs to be remedied. Until we can quantify the abilities of the statistical models to detect different strengths of interactions, we will be kept in the dark about how many unexpected interactions we are missing, and the amount of data required to uncover them.

## Acknowledgments

This research was supported by the United Kingdom Natural Environment Research Council (NERC) via the studentship grant NE/M010481/1, and the research grant NE/V001396/1. The authors declare no conflicts of interest.

## Author contributions

BB and DM performed the analyses and drafted the manuscript. BB derived the critical effect size for the additive null model. DM designed and wrote the code to estimate power to detect non-null interactions. All the authors contributed significantly to the intellectual core of the manuscript; to the interpretation of the results; and to revisions of the manuscript.

## Data availability

All data analysed within this paper is openly available. Code to generate Figure 3 is provided in the Supporting Material.

## Code availability

R code to estimate power, as used to generate Figure 5 can be found at https://github.com/djmurrell/Stressor-Interaction-statistical-power-function.

## References

Ban, S. S., Graham, N. A., and Connolly, S. R. 2014. Evidence for multiple stressor interactions and effects on coral reefs. Global Change Biology, 20(3), 681–697.

Beauchesne, D., Cazelles, K., Archambault, P., Dee, L., and Gravel, D. 2021. On the sensitivity of food webs to multiple stressors. Ecology Letters, 24, 2219–2237. https://doi.org/10.1111/ele.13841

Beaumelle, L., De Laender, F., and Eisenhauer, N. 2020. Biodiversity mediates the effects of stressors but not nutrients on litter decomposition. Elife, 9, e55659.

Bird, G., Wilson, A. E., Williams, G.R., Hardy, N.B. 2021. Parasites and pesticides act antagonistically on honey bee health. Journal of Applied Ecology, 58: 997–1005. https://doi.org/10.1111/1365-2664.13811

Birk, S., Chapman, D., Carvalho, L. et al. 2020. Impacts of multiple stressors on freshwater biota across spatial scales and ecosystems. Nature Ecology and Evolution, 4, 1060–1068 https://doi.org/10.1038/s41559-020-1216-4.

Boone, M.D., Bridges, C.M., Fairchild, J.F. and Little, E.E. 2005. Multiple sublethal chemicals negatively affect tadpoles of the green frog, Rana clamitans. Environmental Toxicology and Chemistry, 24: 1267–1272. https://doi.org/10.1897/04-319R.1

Borenstein, M., Cooper, H., Hedges, L. and Valentine, J., 2009. Effect sizes for continuous data. The Handbook of Research Synthesis and Meta-Analysis, 2, pp.221–235.

Boyd, P. W., Collins, S., Dupont, S., Fabricius, K., Gattuso, J. P., Havenhand, J., and Pörtner, H. O. 2018. Experimental strategies to assess the biological ramifications of multiple drivers of global ocean change—a revie">. Global Change Biology, 24(6), 2239–2261.

Brown, C.J., Saunders, M.I., Possingham, H.P. and Richardson, A.J., 2013. Managing for interactions between local and global stressors of ecosystems. PLoS One, 8(6), p.e65765.

Burgess B. J. and Murrell D. J., 2021. multiplestressR: Additive and Multiplicative Null Models for Multiple Stressor Data. R package version 0.1.1. https://CRAN.R-project.org/package=multiplestressR.

Burgess B. J. and Murrell D. J., 2022. multiplestressR: An R package to analyse factorial multiple stressor data using the additive and multiplicative null models. bioRxiv 2022.04.08.487622; doi: https://doi.org/10.1101/2022.04.08.487622

Burgess, B.J., Purves, D., Mace, G. and Murrell, D.J., 2021. Classifying ecosystem stressor interactions: Theory highlights the data limitations of the additive null model and the difficulty in revealing ecological surprises. Global Change Biology, 27: 3052–3065. https://doi.org/10.1111/gcb.15630

Christensen, M.R., Graham, M.D., Vinebrooke, R.D., Findlay, D.L., Paterson, M.J. and Turner, M.A., 2006. Multiple anthropogenic stressors cause ecological surprises in boreal lakes. Global Change Biology, 12(12), pp.2316–2322.

Cohen, J. (1988). Statistical Power Analysis for the Behavioral Sciences (2nd ed.). Routledge. https://doi.org/10.4324/9780203771587.

Côté, I.M., Darling, E.S. and Brown, C.J., 2016. Interactions among ecosystem stressors and their importance in conservation. Proceedings of the Royal Society B: Biological Sciences, 283(1824), p.20152592.

Crain, C.M., Kroeker, K. and Halpern, B.S., 2008. Interactive and cumulative effects of multiple human stressors in marine systems. Ecology Letters, 11(12), pp.1304–1315.

De Laender, F., 2018. Community-and ecosystem-level effects of multiple environmental change drivers: Beyond null model testing. Global Change Biology, 24(11), pp.5021–5030.

Dey, C.J. and Koops, M.A., 2021. The consequences of null model selection for predicting mortality from multiple stressors. Proceedings of the Royal Society B, 288(1948), p.20203126.

Dixon, G., Abbott, E., and Matz, M. 2020. Meta-analysis of the coral environmental stress response: Acropora corals show opposing responses depending on stress intensity. Molecular Ecology, 29(15), 2855–2870.

Filazzola, A., and Cahill, J. F. 2021. Replication in field ecology: Identifying challenges and proposing solutions. Methods in Ecology and Evolution, 12, 1780–1792. https://doi.org/10.1111/2041-210X.13657

Flügge, A.J., Olhede, S.C. and Murrell, D.J. 2012. The memory of spatial patterns: changes in local abundance and aggregation in a tropical forest. Ecology, 93: 1540–1549.

Fournier, V., Rosenheim, J.A., Brodeur, J., Diez, J.M. and Johnson, M.W., 2006. Multiple plant exploiters on a shared host: testing for nonadditive effects on plant performance. Ecological Applications, 16(6), pp.2382–2398.

Gomez Isaza, D.F., Cramp, R.L. and Franklin, C.E., 2020. Living in polluted waters: a meta-analysis of the effects of nitrate and interactions with other environmental stressors on freshwater taxa. Environmental Pollution, p.114091.

Gurevitch, J., Morrison, J.A. and Hedges, L.V., 2000. The interaction between competition and predation: a meta-analysis of field experiments. The American Naturalist, 155(4), pp.435–453.

Halpern, B. S., Frazier, M., Potapenko, J., Casey, K. S., Koenig, K., Longo, C., and Walbridge, S.s 2015. Spatial and temporal changes in cumulative human impacts on the world’s ocean. Nature Communications, 6(1), 1–7.

Hanson, P. J., and Walker, A. P 2020. Advancing global change biology through experimental manipulations: Where have we been and where might we go? Global Change Biology, 26(1), 287–299.

Hedges, L. V., and I. Olkin., 1985. Statistical methods for meta-analysis. Academic Press, New York.

Hering, D., Carvalho, L., Argillier, C., Beklioglu, M., Borja, A., Cardoso, A. C., … and Birk, S 2015. Managing aquatic ecosystems and water resources under multiple stress—An introduction to the MARS project. Science of the Total Environment, 503, 10–21.

Hodgson, E.E. and Halpern, B.S., 2018. Investigating cumulative effects across ecological scales. Conservation Biology, 33(1), pp.22–32.

Jackson, M.C., Loewen, C.J., Vinebrooke, R.D. and Chimimba, C.T., 2016. Net effects of multiple stressors in freshwater ecosystems: a meta-analysis. Global Change Biology, 22(1), pp.180–189.

Jackson, M.C., Pawar, S. and Woodward, G., 2021. The Temporal Dynamics of Multiple Stressor Effects: From Individuals to Ecosystems. Trends in Ecology and Evolution, 36(5): 402–410.

Johnson, P.C., Barry, S.J., Ferguson, H.M. and Müller, P., 2015. Power analysis for generalized linear mixed models in ecology and evolution. Methods in Ecology and Evolution, 6(2), pp.133–142.

Lafferty, K.D. and Holt, R.D. 2003. How should environmental stress affect the population dynamics of disease? Ecology Letters, 6: 654–664. https://doi.org/10.1046/j.1461-0248.2003.00480.x

Lajeunesse, M.J., 2011. On the meta-analysis of response ratios for studies with correlated and multi-group designs. Ecology, 92(11), pp.2049–2055.

Lakens, D. 2017. Equivalence Tests: A Practical Primer for t Tests, Correlations, and Meta-Analyses. Social Psychological and Personality Science, 8(4): 355-362. doi:10.1177/1948550617697177

Lakens, D. 2022. Sample Size Justification. Collabra: Psychology, 8 (1): 33267. doi: https://doi.org/10.1525/collabra.33267

Lange, K., Bruder, A., Matthaei, C.D., Brodersen, J. and Paterson, R.A., 2018. Multiple-stressor effects on freshwater fish: Importance of taxonomy and life stage. Fish and Fisheries, 19(6), pp.974–983.

Lemm, J. U., Venohr, M., Globevnik, L., Stefanidis, K., Panagopoulos, Y., van Gils, J., and Birk, S. 2021. Multiple stressors determine river ecological status at the European scale: Towards an integrated understanding of river status deterioration. Global Change Biology, 27(9), 1962–1975.

Ma, Z., Chen, H. Y., Li, Y., and Chang, S. X 2020. Interactive effects of global change factors on terrestrial net primary productivity are treatment length and intensity dependent. Journal of Ecology, 108(5), 2083–2094.

Maltby, L. 1999. Studying stress: The importance of organism-level responses. Ecological Applications, 9(2), 431–440.

Mudge, J.F., Baker, L.F., Edge, C.B. and Houlahan, J.E., 2012. Setting an optimal α that minimizes errors in null hypothesis significance tests. PLoS One, 7(2), p.e32734.

Murrell, D.J 2018. A global envelope test to detect non-random bursts of trait evolution. Methods in Ecology and Evolution, 9: 1739–1748.

Nõges, P., Argillier, C., Borja Á., Garmendia, J.M., Hanganu, J., Kodeš, V., Pletterbauer, F., Sagouis, A., Birk, S. 2016. Quantified biotic and abiotic responses to multiple stress in freshwater, marine and ground waters. Science of the Total Environment. 540: 43-52. doi: 10.1016/j.scitotenv.2015.06.045. Epub 2015

Orr, J.A., Vinebrooke, R.D., Jackson, M.C., Kroeker, K.J., Kordas, R.L., Mantyka-Pringle, C., Van den Brink, P.J., De Laender, F., Stoks, R., Holmstrup, M. and Matthaei, C.D., 2020. Towards a unified study of multiple stressors: divisions and common goals across research disciplines. Proceedings of the Royal Society B, 287(1926), p.20200421.

Polazzo, F., Roth, S. K., Hermann, M., Mangold-Döring, A., Rico, A., Sobek, A., and Jackson, M. C. 2021. Combined effects of heatwaves and micropollutants on freshwater ecosystems: Towards an integrated assessment of extreme events in multiple stressors research. Global Change Biology 00, 1–20. https://doi.org/10.1111/gcb.15971.

Przeslawski, R., Byrne, M., and Mellin, C 2015. A review and meta-analysis of the effects of multiple abiotic stressors on marine embryos and larvae. Global Change Biology, 21(6), 2122–2140.

Rajala, T., Olhede, S.C., Murrell, D.J. 2019. When do we have the power to detect biological interactions in spatial point patterns? Journal of Ecology, 107: 711–721.

Rineau, F., Malina, R., Beenaerts, N., Arnauts, N., Bardgett, R. D., Berg, M. P., and Vangronsveld, J. 2019. Towards more predictive and interdisciplinary climate change ecosystem experiments. Nature Climate Change, 9(11), 809–816.

Schäfer, R.B. and Piggott, J.J., 2018. Advancing understanding and prediction in multiple stressor research through a mechanistic basis for null models. Global Change Biology, 24(5), pp.1817–1826.

Seifert, M., Rost, B., Trimborn, S., and Hauck, J. 2020. Meta-analysis of multiple driver effects on marine phytoplankton highlights modulating role of pCO2. Global Change Biology, 26(12), 6787–6804.

Senior, A. M., Grueber, C. E., Kamiya, T., Lagisz, M., O’dwyer, K., Santos, E. S., and Nakagawa, S. 2016. Heterogeneity in ecological and evolutionary meta-analyses: its magnitude and implications. Ecology, 97(12), 3293–3299.

Simmons, B.I., Blyth, P.S.A., Blanchard, J.L. et al. 2021. Refocusing multiple stressor research around the targets and scales of ecological impacts. Nature Ecology and Evolution, 5: 1478–1489 https://doi.org/10.1038/s41559-021-01547-4.

Siviter, H., Bailes, E. J., Martin, C. D., Oliver, T. R., Koricheva, J., Leadbeater, E., and Brown, M. J. 2021. Agrochemicals interact synergistically to increase bee mortality. Nature, 596(7872), 389–392.

Sokolova, I., 2021. Bioenergetics in environmental adaptation and stress tolerance of aquatic ectotherms: linking physiology and ecology in a multi-stressor landscape. Journal of Experimental Biology, 224(Suppl_1). doi:10.1242/jeb.236802.

Spears, B. M., Chapman, D. S., Carvalho, L., Feld, C. K., Gessner, M. O., Piggott, J. J., and Birk, S. 2021. Making waves. Bridging theory and practice towards multiple stressor management in freshwater ecosystems. water Research, 116981.

Timmers, M.A.,Jury, C.P., Vicente, J., Bahr, K.D., Webb, M., and Toonen, R.J. 2021 Biodiversity of coral reef cryptobiota shuffles but does not decline under the combined stressors of ocean warming and acidification. Proceedings of the National Academy of Sciences, 118 (39) e21032751.

van Veen, F.J.F. and Murrell, D.J., 2005. A simple explanation for universal scaling relations in food webs. Ecology, 86(12), pp.3258–3263

Viechtbauer, W. 2010. Conducting meta-analyses in R with the metafor package. Journal of Statistical Software, 36(3), 1–48. https://doi.org/10.18637/jss.v036.i03

Yang, Y., Hillebrand, H., Lagisz, M., Cleasby, I., and Nakagawa, S. 2022. Low statistical power and overestimated anthropogenic impacts, exacerbated by publication bias, dominate field studies in global change biology. Global Change Biology, 28, 969–989. https://doi.org/10.1111/gcb.15972.

