## Supplementary Information for "Are experiment sample sizes adequate to detect biologically important interactions between multiple stressors?"

***Additive null model***

Here we give the full details of the additive null model outlined by Gurevitch et al., (2000). The additive effect size (${ES}_{Add})$is defined as:

$${ES}_{Add}=\frac{\bar{X}_{I}-\bar{X}_{A}-\bar{X}_{B}+\bar{X}_{C}}{s}\cdot J$$

$$S1.1)$$

where the pooled standard deviation, *s*, is defined as:

$$s= \sqrt{\frac{\left( N_{I}-1 \right)\cdot\left( {SD}_{I} \right)^{2} + \left( N_{A}-1 \right)\cdot\left( {SD}_{A} \right)^{2} + \left( N_{B}-1 \right)\cdot\left( {SD}_{B} \right)^{2} + \left( N_{C}-1 \right)\cdot\left( {SD}_{C} \right)^{2}}{N_{I}+ N_{A}+ N_{B}+N_{C}-4}}$$

$$S1.4)$$

and the small sample size bias correction factor is defined as:

$$J = 1- \frac{3}{4\cdot\left( N_{I}+ N_{A}+ N_{B}+N_{C}-4 \right)-1}$$

$$S1.4)$$

$J$is used to account for the inherent bias in effect size estimation when sample sizes are small (Borenstein et al., 2009).

The variance of ${ES}_{Add}$is defined as:

$V_{Add} = J^{2}\cdot\left[ \frac{1}{N_{I}}+\frac{1}{N_{A}}+\frac{1}{N_{B}}+\frac{1}{N_{C}}+\frac{\left( {ES}_{Add} \right)^{2}}{2\left( N_{I}+ N_{A}+ N_{B}+ N_{C} \right)} \right]$

$$S1.5)$$

The variance is used to estimate the standard error as:

$${SE}_{Add} = \sqrt{V_{Add}}$$

$$S1.6)$$

The confidence intervals are computed using the standard error in the normal way:

$${CI}_{Add} = Z_{\alpha/2}\cdot{SE}_{Add}$$

$$S1.7)$$

with $Z_{\alpha/2}$ being the critical Z-score taken at the statistical level of significance *α.* Typically, *α* = 0.05, and we divide by two as a two-tailed test is required because the stressors interaction can be less than, or greater than expected under the null model, which means $Z_{\alpha/2}=1.96$.

***Multiplicative null model***

In addition to the additive null model, we implement the form of the multiplicative null model detailed by Lajeunesse (2011). The multiplicative effect size, (*ES_Mul_*) is defined as

${ES}_{Mul} = ln \left( \bar{X}_{I} \right)- \ln(\bar{X}_{A})-\ln(\bar{X}_{B})+ \ln(\bar{X}_{C})$.

$$S2.1)$$

The corresponding variance, (*V_Mul_*), is defined as

$V_{Mul}= \frac{{({SD}_{I})}^{2}}{{(\bar{X}_{I})}^{2}\cdot N_{I}}+\frac{{({SD}_{A})}^{2}}{{(\bar{X}_{A})}^{2}\cdot N_{A}}+\frac{{({SD}_{B})}^{2}}{{(\bar{X}_{B})}^{2}\cdot N_{B}}+\frac{{({SD}_{C})}^{2}}{{(\bar{X}_{C})}^{2}\cdot N_{C}}$.

$$S2.2)$$

This leads to the standard error, (*SE_Mul_*), being computed as

${SE}_{Mul}= \sqrt{V_{Mul}}$,

$$S2.3)$$

and the confidence intervals (*CI_Mul_*), are determined as

${CI}_{Mul} = Z_{\alpha/2}\cdot{SE}_{Mul}$,

$$S2.4)$$

where, as above, we set the significance level to 0.05 (Z_α/2_ = 1.96), hence 95% confidence intervals are calculated.

We draw the reader’s attention to the observation that, in contrast to the additive null model (Equation S1) the variance for the multiplicative model (Equation S2.2) is independent of the effects size.

***Interaction classifications***

As described in the main text interaction can be classified as a synergistic, antagonistic, reversal, or null interaction using either the additive (Gurevitch et al., 2000) or multiplicative (Lajeunesse, 2011) null models. The classification of an interaction is dependent upon the effect size ($ES$), associated confidence intervals (${CI}_{95\%}$), the observed $X_{O}$ and expected $X_{E}$ interactive effect of stressors.

For the additive null model, the $X_{E}$ can be calculated as $\bar{X}_{A}+\bar{X}_{B}-2\bar{X}_{C}$; while for the multiplicative null model, $X_{E}$ can be calculated as $\ln\left( \frac{\bar{X}_{A}\bar{X}_{B}}{\bar{X}_{C}^{2}} \right)$. Similarly, for the additive null model, $X_{O}$ can be calculated as $\bar{X}_{I}-\bar{X}_{C}$; while for the multiplicative null model, $X_{O}$ Can be calculated as $\ln\left( \frac{\bar{X}_{I}}{\bar{X}_{C}} \right)$. Using this scheme, interactions were classified as follows:

When $X_{E}$ ≥ 0

1) If $ES \pm{CI}_{95\%}$ overlaps zero, then the interaction was assigned a null classification.
2) If $ES$ > 0, the interaction was assigned a synergistic classification.
3) If $ES$ < 0, and if$X_{O}$ and $X_{E}$both have the same polarity, the interaction was assigned an antagonistic classification.
4) If $ES$ < 0, and if$X_{O}$ and ­$X_{E}$ have contrasting polarities, the interaction was assigned a reversal classification.

When $X_{E}$< 0

1) If $ES \pm{CI}_{95\%}$ overlaps zero, then the interaction was assigned a null classification.
2) If $ES$ < 0, the interaction was assigned a synergistic classification.
3) If $ES$ > 0, and if$X_{O}$ and $X_{E}$ both have the same polarity, the interaction was assigned an antagonistic classification.
4) If $ES$ > 0, and if$X_{O}$ and $X_{E}$ have contrasting polarities, the interaction was assigned a reversal classification.

***Critical Effect Size***

By combining and rearranging Equations S1.1 – S1.6, it is possible to express the additive effect size as a function of sample size:

$$\left| {ES}_{Add} \right|> \sqrt{\left( \frac{1}{N_{C}}+ \frac{1}{N_{A}}+ \frac{1}{N_{B}}+ \frac{1}{N_{I}} \right)\cdot\left( \frac{2\cdot{(Z_{\alpha/2})}^{2}\cdot\left( N_{C}+ N_{A}+ N_{B}+ N_{I} \right)\cdot J^{2}}{2\cdot\left( N_{C}+ N_{A}+ N_{B}+ N_{I} \right)- {(Z_{\alpha/2})}^{2} \cdot J^{2}} \right)}$$

*(Inequality S1)*

This inequality shows that for a given sample size, there is a minimum additive effect size value (also been referred to as the critical effect size by Lakens 2022) which must be exceeded before the interaction can be correctly classified as being non-null.

Unfortunately, due to a more complex structure in the calculation of the variance the multiplicative null model does not yield such an inequality for calculating the critical effect size. However, we repeat the data analysis shown in Figure 3, but using the multiplicative null model instead of the additive null model (Figure S1). Although the relationship is much less straightforward there appears to be a higher probability of rejecting the null model expectation in favour of one of the three alternative interactions (antagonistic, reversal, synergistic).

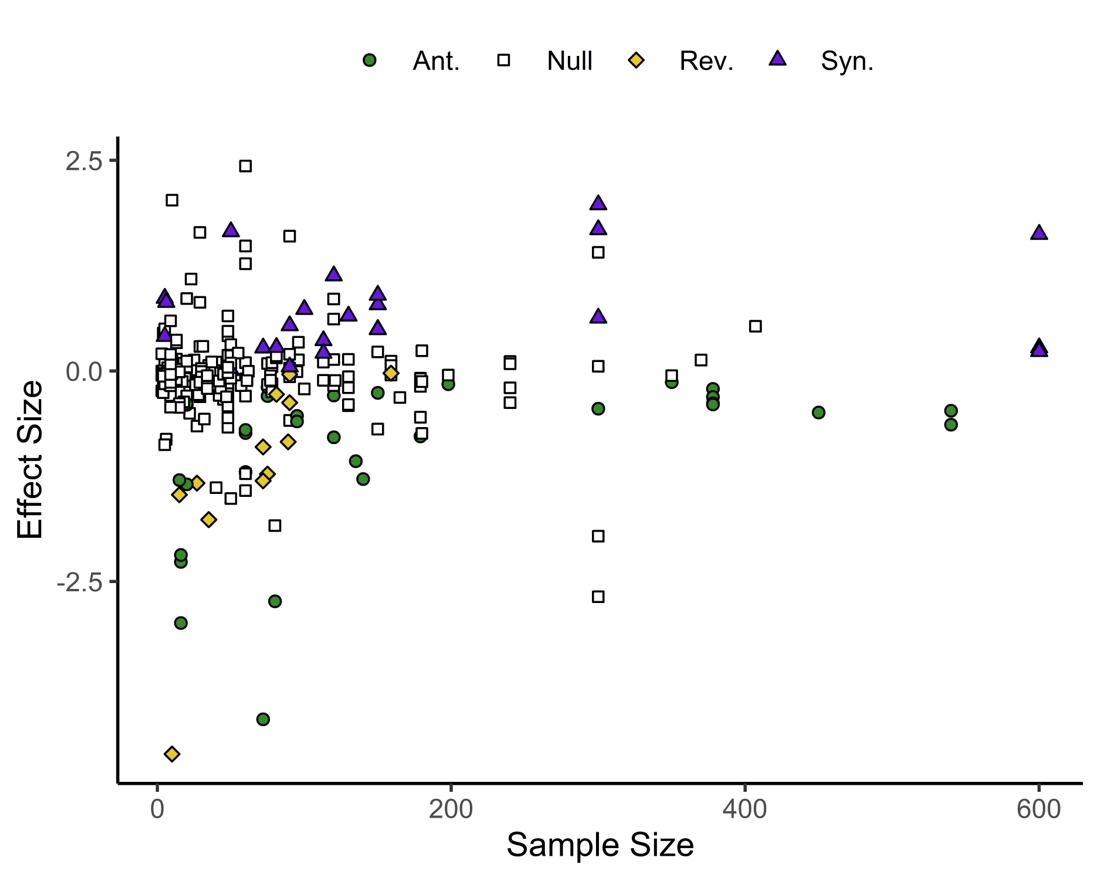

**Figure S1.** The effect of sample size on the ability to detect interactions with different effect sizes for the bee health responses to multiple stressors in Siviter et al. (2021). Open squares denote data points that are statistically indistinguishable from the null model of a multiplicative interaction (Equation S2). Data points that lead to the rejection of the null model can be assigned as synergistic (blue triangles), antagonistic (green circles), or reversals (orange diamond). Median sample size per treatment is plotted on the x axis. This figure was generated using the *multiplestressR* R package (Burgess & Murrell, 2022).

**Statistical power**

Here we show how changing the standard deviation of the treatment values has a negligible effect on power for the additive null model (Figure S2), and repeat the analysis on statistical power shown in Figure 5a of the main text, but this time using the multiplicative null model (Figure S3). The reader should recall that the additive and multiplciative null models make different assumptions about the nature of the stressor interactions and that they can therefore return different types of interaction for the same dataset. Hence in Figure S3 we find a weakly positive effect size implying a synergy even though interactions are all antagonistic under the additive null model (Figure 5a).

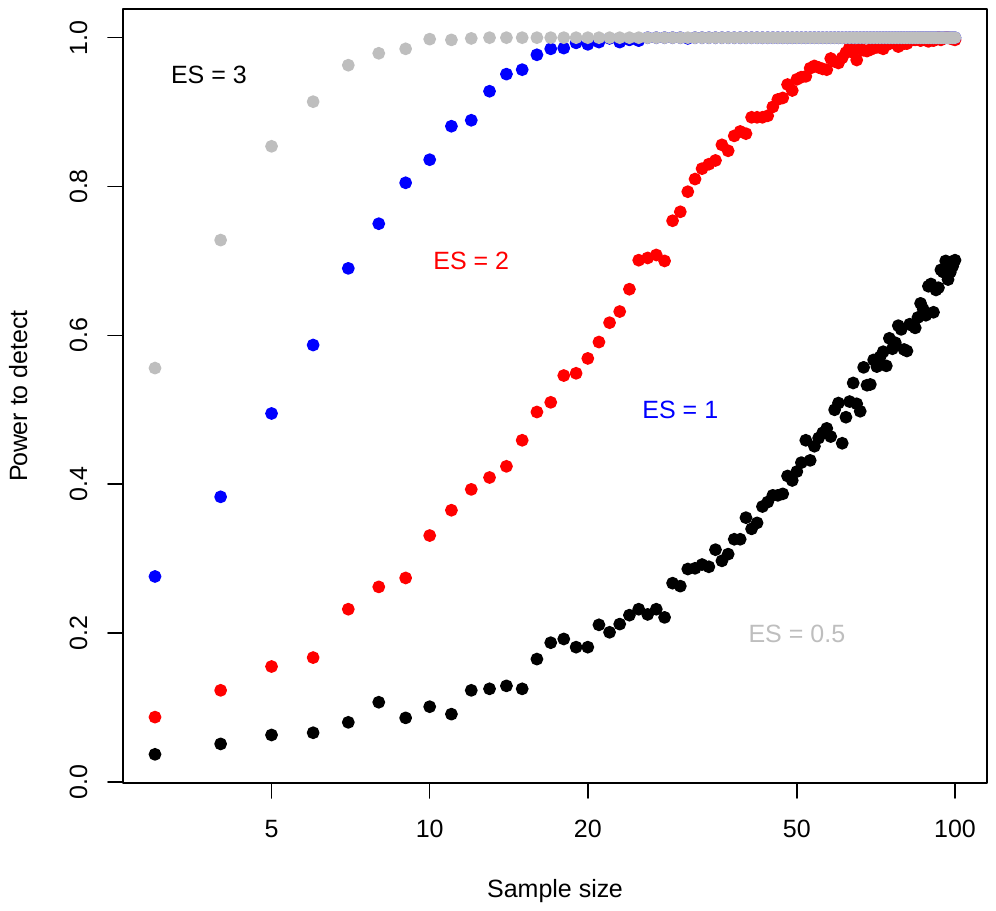

**Figure S2.** Treatment variation has little effect on statistical power in the additive null model when holding effect sizes constant. Expected effect sizes (ES) are the same as in Figure 5a of the main text, but now we assume the standard deviation of the treatment variables are smaller, being equal to $\sigma_{x}=0.025$. This necessitates selecting expected mean interaction treatment values of ${E(\bar{X}}_{I})=\in\left\{ 0.5125, 0.525, 0.55, 0.575 \right\}$ in order to generate effect sizes ${ES}_{ADD}=\{3, 2, 1, 0.5\}$ respectively. All other data simulation parameters are as described in the main text for Figure 5.

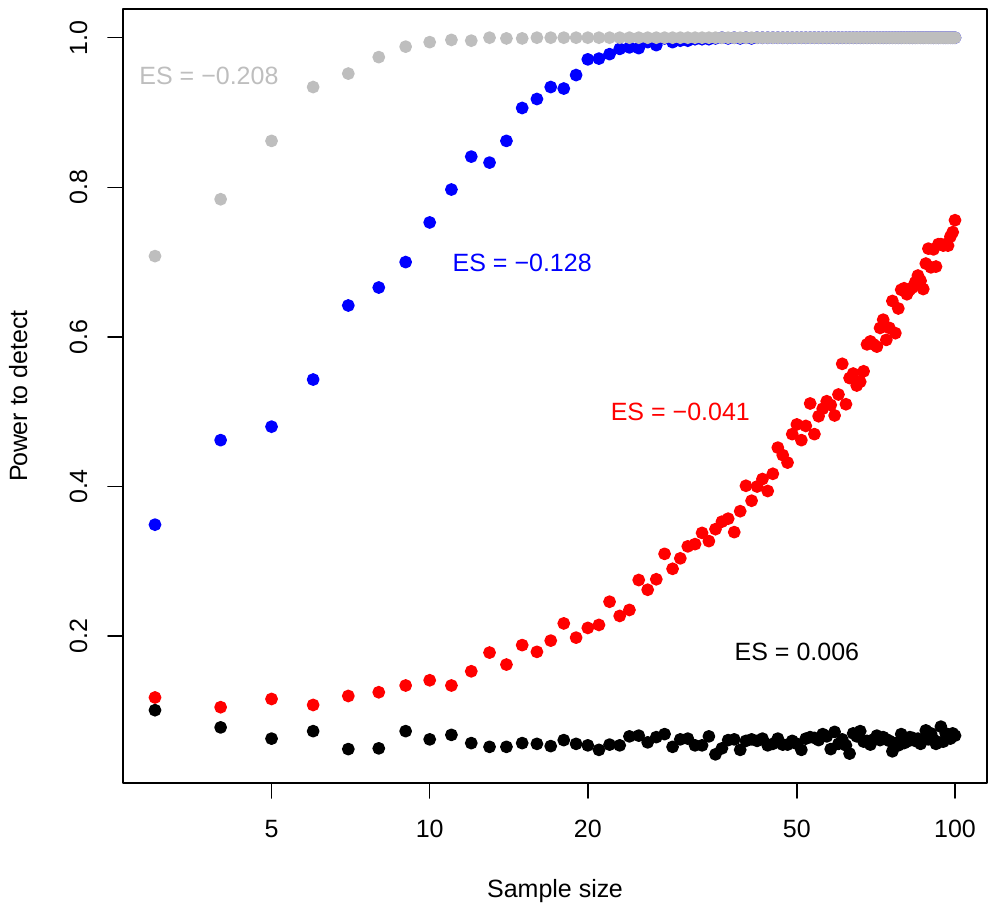

**Figure S3.** The effect of sample size on statistical power for the multiplicative null model (equations S1). Data is generated using the simulation method described in the main text. Parameters used in the simulation are as for Figure 5a in the main text. We draw the reader’s attention to the fact that the effect sizes ${ES}_{Mul}$ are on a different scale than for the additive model and that interactions (the sign of the effect sizes) may be different between the two models.

**Code to produce power estimates**

The code we use to generate Figure 5 and Figure S3 is freely available and can be found at https://github.com/djmurrell/Stressor-Interaction-statistical-power-function. Users should note that the code assumes data are distributed according to a Gaussian distribution. This might be appropriate for many types of response (e.g., density), but other responses may be better described by a different distribution. The code can be adapted for such non-Gaussian data. The code is written to implement either additive or multiplicative nulls models, but we caution the user that, in its current form, the code does not verify the predicted interaction for the additive model returns ‘sensible’ values (e.g., is positive, and in the case of survival/mortality rates is not above 1).

**References**

Borenstein, M., Cooper, H., Hedges, L. and Valentine, J., 2009. Effect sizes for continuous data. *The Handbook of Research Synthesis and Meta-Analysis*, 2, pp.221-235.

Burgess, B. J., & Murrell, D. J. (2022). *multiplestressR*: An R package to analyse factorial multiple stressor data using the additive and multiplicative null models. *bioRxiv*.

Gurevitch, J., Morrison, J.A. and Hedges, L.V., 2000. The interaction between competition and predation: a meta-analysis of field experiments. *The American Naturalist*, 155(4), pp.435-453.

Lajeunesse, M.J., 2011. On the meta‐analysis of response ratios for studies with correlated and multi‐group designs. *Ecology*, 92(11), pp.2049-2055.

Lakens, D. 2022. Sample Size Justification. *Collabra: Psychology*, 8 (1): 33267. doi: <https://doi.org/10.1525/collabra.33267>

Siviter, H., Bailes, E. J., Martin, C. D., Oliver, T. R., Koricheva, J., Leadbeater, E., & Brown, M. J. 2021. Agrochemicals interact synergistically to increase bee mortality. *Nature*, 596(7872), 389-392.
